# Construction of *lux*-based promoter-reporter platforms in *Mycobacterium bovis* BCG for screening of drug repurposing small-molecule compounds as new anti-tuberculosis drugs

**DOI:** 10.1101/2021.11.05.467536

**Authors:** Li Zhu, Annie Wing-Tung Lee, Kelvin Ka-Lok Wu, Peng Gao, Kingsley King-Gee Tam, Rahim Rajwani, Galata Chala Chaburte, Timothy Ting-Leung Ng, Chloe Toi-Mei Chan, Hiu Yin Lao, Wing Cheong Yam, Richard Yi-Tsun Kao, Gilman Kit Hang Siu

## Abstract

The emergence of multidrug-resistant strains and hyper-virulent strains of *Mycobacterium tuberculosis* are big therapeutic challenges for tuberculosis (TB) control. Repurposing bioactive small-molecule compounds has recently become a new therapeutic approach against TB. This study aimed to construct a rapid screening system to identify novel anti-TB agents from a library of small-molecule compounds.

In this study, a total of 320 small-molecule compounds were used to screen for their ability to suppress the expression of a key virulence gene, *phoP*, of *M. tuberculosis* complex using luminescence (*lux*)-based promoter-reporter platforms. The minimum inhibitory and bactericidal concentrations on drug-resistant *M. tuberculosis* and cytotoxicity to human macrophage were determined. RNA-sequencing (RNA-seq) was conducted to determine the drug mechanisms of the selected compounds as novel antibiotics or anti-virulent agents against the *M. tuberculosis* complex.

Six compounds displayed bactericidal activity against *M. bovis* BCG, in which Ebselen demonstrated the lowest cytotoxicity to macrophage and was considered as a potential antibiotic for TB. Another ten compounds did not inhibit the *in vitro* growth of the *M. tuberculosis* complex but down-regulated the expression of *phoP* specifically. Of them, ST-193 and ST-193 (hydrochloride) showed low cytotoxicity and could dysregulate the entire *phoP*-associated gene network, and thus identified as potential anti-virulence agents for *M. tuberculosis*. This study provides a rapid screening platform coupled with a systematic validation and eventually suggested one potential antibiotic and two anti-virulence agents for *M. tuberculosis* infections.

## 1. Introduction

Tuberculosis (TB) remains a significant global health problem today. According to the World Health Organization Global TB report in 2020, TB ranked tenth among the leading causes of death worldwide [1]. Approximately 10 million people developed TB, which claimed over 1.4 million lives in 2019 [1]. TB infections associated with multidrug-resistant (MDR), extensively drug-resistant (XDR), and totally drug-resistant (TDR) *Mycobacterium tuberculosis* strains have been increasing in recent years [1–4]. This situation poses a significant threat to global TB control, particularly in resource-poor countries with a high prevalence of AIDS. However, according to the global new TB drug development pipeline built by the Stop TB Partnership Working Group on New Drugs in 2017, only four anti-TB drugs were in phase III clinical trials [5], and some of them could not penetrate the host cell, thereby failing to eliminate *M. tuberculosis* [6]. Hence, there is an urgent need for developing new anti-TB treatment against *M. tuberculosis* strains with various patterns of drug resistance.

Anti-virulence therapy is an innovative therapeutic strategy to treat multidrug-resistant organisms. It focuses on disarming bacterial virulence factors that facilitate disease development instead of killing the bacteria [7]. A previous study discovered a small-molecule compound, 2-phospho-L-ascorbic, could reduce mycobacterial survival in macrophage infections [8]. It underlined the potential for the development of anti-virulence agents against *M. tuberculosis. PhoP* is a global transcriptional regulator of lipid metabolism and hypoxic response and controls the expression of ~2% of the genes in *M. tuberculosis* [9]. It was shown that disruption of *phoP* in *M. tuberculosis* caused impaired multiplication within macrophages, suggesting that this gene possibly plays an essential role in the intracellular growth of *M. tuberculosis* [10]. In addition, our previous study identified a common mutation in the promoter region of *phoP*, which could confer aggressive intramacrophage growth of hypervirulent *M. tuberculosis* strains [11]. Therefore, we hypothesized that suppression of *phoP* expression might be able to impair intramacrophage survival of *M. tuberculosis*, facilitating the host’s immune system to eradicate the bacteria.

Over thousands of small-molecule compounds are approved and passing phase I clinical drugs, meaning that they have completed extensive preclinical and clinical studies and have well-characterized bioactivities, safety, and bioavailability properties. These compounds could be potentially repurposed to inhibit *M. tuberculosis* virulence. As such, a rapid and high-throughput platform that can screen effective anti-virulence agents for TB is warranted.

The present study aimed to develop a *lux*-based promoter-reporter platform for massive screening of small-molecule compounds that were likely to suppress the expression of *phoP* in *Mycobacterium bovis* BCG. Compared with *M. tuberculosis, M. bovis* BCG has a lower biosafety level (Risk Group 2, Biosafety Level 2 Practices), despite its genome being>99.95% identical to that of *M. tuberculosis* reference strain H37Rv [12]. This allowed the screening process to be done in the BSL-2 laboratory. A library of 320 small-molecule compounds with antiviral activities was screened for their anti-TB potency. The compounds that could reduce the *lux* signal were selected for further validation on (i) their abilities to inhibit *in vitro* growth of *M. tuberculosis* complex, (ii) cytotoxicity to THP-1 macrophage, and (iii) dysregulated expression of *phoP* and its associated gene networks. Our findings eventually suggested one potential antibiotic and two anti-virulence agents for anti-TB therapy.

## 2. Materials and Methods

### 2.1. Bacterial strains

*Mycobacterium bovis* vaccine strain, Bacille Calmette-Guérin (BCG)-1 [Russia] (*M. bovis* BCG), which was used for high-throughput screening of small compounds, was collected from Queen Mary Hospital, Hong Kong. The organism was revived using Middlebrook 7H9 broth and 7H10 agar including OADC (10%), glycerol (0.2% in 7H9 broth and 0.5% in 7H10 agar), sodium pyruvate (4.4mg/ml), and pancreatic digest of casein (1mg/ml) at 37°C for 14 days.

*M. tuberculosis* H37Rv and two drug-resistant clinical isolates of *M. tuberculosis*, namely HKU1462 (an MDR-TB strain co-resistant to isoniazid and rifampicin) and WC274 (an XDR-TB strain resistant to isoniazid, rifampicin, pyrazinamide, ethambutol, streptomycin, fluoroquinolones, and injectable aminoglycoside) collected from the same hospital were used to determine the inhibitory and bactericidal activities against *M. tuberculosis*. Resuscitation of archived clinical isolates was performed using Lowenstein-Jensen (LJ) medium. The detailed drug susceptibility patterns of the clinical isolates were shown in Supplementary Table S1. The Biosafety Committee at the Hong Kong Polytechnic University approved all the experiments (Code: RSA18020).

### 2.2. Construction of *lux*-based promoter-reporter plasmids and a negative control

Mycobacterial reporter plasmid -pMV306G13+Lux was obtained from Brian Robertson and Siouxsie Wiles (Addgene plasmid #26160, Figure 1a) [13], containing *luxCDABE*, a G13 promoter, and a kanamycin-resistant gene. G13 promoter was replaced by *phoP* promoter (Figure 1c, Supplementary Table S2A) to construct pMV306PhoP+Lux. The negative control - pMV306Adaptor+Lux was similarly created by replacing the G13 promoter with an adaptor (Figure 1b, 5’-GGCCGCTTAGATCTTTC-3’- a random sequence without the promoter activity). All plasmids used in this study are listed in Supplementary Table S2B.

**Fig. 1.**
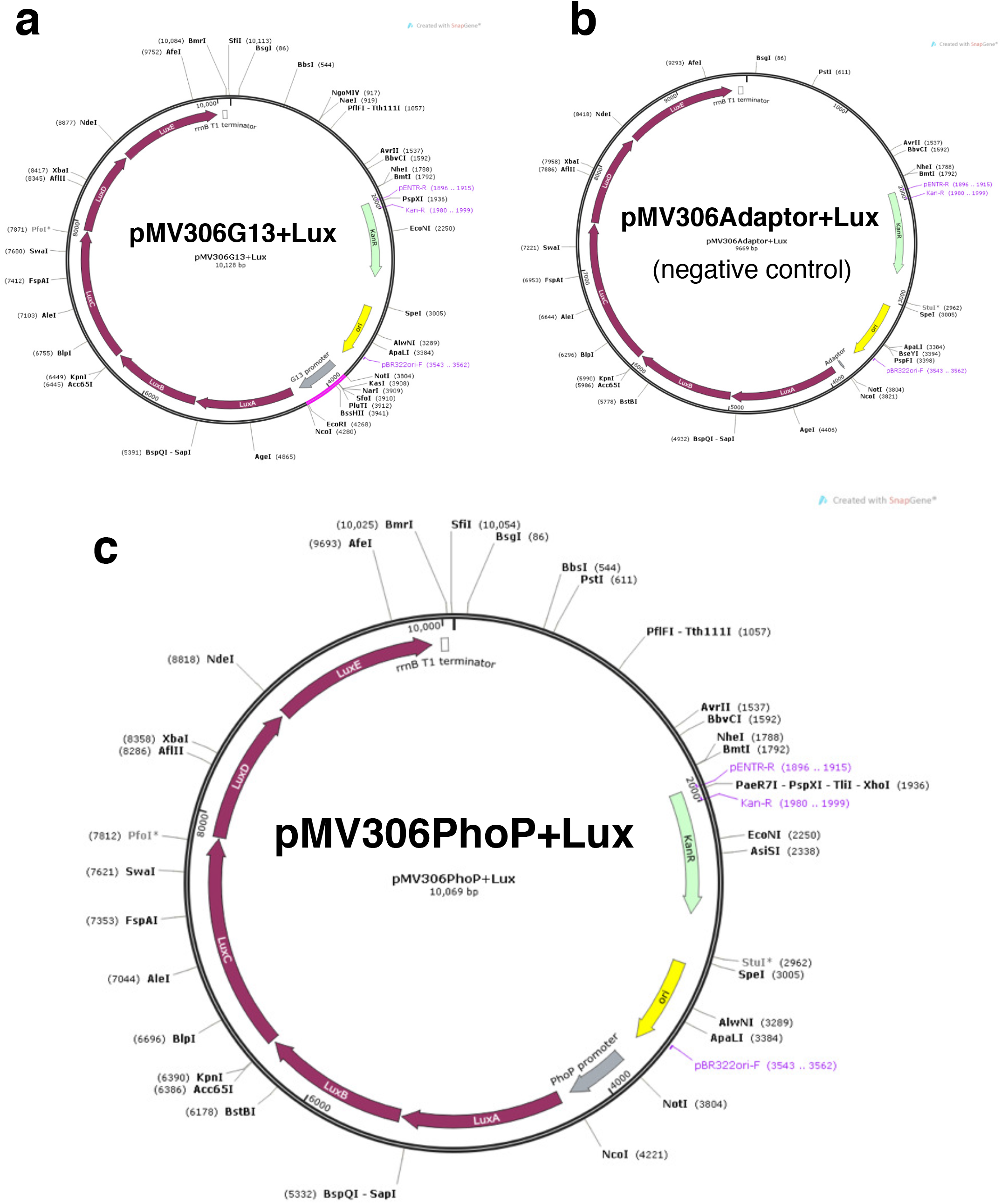
Construction of the *lux*-based promoter-reporter platform. **a,** pMV306G13+Lux **b,** pMV306Adaptor+Lux (negative control) and **c,** pMV306PhoP+Lux.

### 2.3. Transformation of *lux*-based reporter plasmids into *M. bovis* BCG

*Lux*-based promoter-reporter plasmids (pMV306PhoP+Lux) and negative control (pMV306Adaptor+Lux) were transformed into *M. bovis* BCG by electroporation. The sequences of the plasmid were confirmed by PCR-sequencing. The *lux* signal of transformed *M. bovis* BCG was validated by an IVIS Lumina imaging system (Perkin-Elmer, China).

### 2.4. Validating the correlation of *lux* signal of the promoter-reporter screening platform with *phoP* gene expression in *M. bovis* BCG

Ethoxzolamide (ETZ) was shown to inhibit PhoPR regulon in *Mycobacterium tuberculosis* via binding to *phoP* promoter regions [14]. In this study, *M. bovis* BCG transformed with pMV306PhoP+Lux were treated with 200μg/ml ethoxzolamide (experimental group) and DMSO (control group) at 37°C for 24 hours. Optical Density via absorbance at 600 nm (OD_600_) and *lux* signals from all samples were measured by Benchmark Plus Microplate Spectrophotometer (BIO-RAD) and VICTOR3 Multilabel Plate Reader (PerkinElmer), respectively. Total RNA was then extracted from the experimental and control group for quantitative reverse transcription PCR (RT-qPCR). The fold-changes in gene expression of *luxA* and *phoP* were calculated based on a housekeeping gene *recX* [15]. The primers used for RT-qPCR are listed in Supplementary Table S2C.

### 2.5. Screening experiments

The screening experiments involved *M. bovis* BCG transformed with pMV306PhoP+Lux and pMV306Adaptor+Lux (negative control). Both transformants were challenged with a total of 320 antiviral small compounds (MedChemExpress, USA) and three first-line anti-TB drugs (ethambutol, isoniazid, and rifampicin) (Supplementary Table S3). Considering bactericidal effects of anti-TB drugs, the final concentrations of three anti-TB drugs in screening were adjusted to two-fold below their respective MICs to *M. bovis* BCG [16]. The small compounds were dissolved in DMSO and water where appropriate. For each group, each compound (2-μl, 10-mM) was dispensed in 100-μl of *M. bovis* BCG suspension (OD_600_≈0.2) in triplicate. We also prepared, in triplicate, 100-μl of BCG suspensions that excluded the compound treatment as drug-free control. The final concentration of the compound library was 200-μM, and the final concentrations of the anti-TB drugs were 3.8μg/ml (ethambutol), 0.08μg/ml (isoniazid), and 0.4μg/ml (rifampicin). OD_600_ and *lux* were measured and compared after 4 hours of treatment with the compounds.

### 2.6. Observation of *M. bovis* BCG survival after treatments of 16 compounds

Sixteen compounds were shown to have total *lux* inhibition (CPS -photon count per second ≈0) after 4-hour treatment. The respective *M. bovis* BCG suspensions were washed with 7H9 medium twice to remove residual compounds. The washed bacterial pellets were then inoculated onto drug-free 7H10 medium, followed by incubating at 37°C for 14 days to determine the viability of the compound-treated *M. bovis* BCG.

### 2.7. Determining minimal inhibitory concentrations (MICs) and minimal bactericidal concentrations (MBCs) against *M. tuberculosis* complex

The MICs and MBCs of the selected compounds against *M. bovis* BCG, *M. tuberculosis* H37Rv, HKU14621 (MDR clinical isolate) and WC276 (XDR clinical isolate) were determined using the microbroth dilution method according to the Clinical and Laboratory Standards Institute (CLSI) [17]. The concentrations for the selected compounds were 2-fold serially diluted from 400-μM to 1.5625-μM.

### 2.8. RNA-Seq transcriptome analysis

For RNA-seq analysis, untransformed *M. bovis* BCG (i.e. with no plasmid) was treated for 4 hours with the selected compounds at a concentration of 200-μM or two-fold lower than their respective MICs if they exhibited inhibitory or bactericidal activity in previous MIC and MBC experiments. Total RNA was extracted and removed rRNA after a 4-hour treatment, followed by a strand-specific library (the bacterial cDNA library) construction with the quality control process. Subsequently, RNA-Seq was performed using the HiSeq X Ten system. Differential mRNA expression was analysed by DESeq2 using Trinity v2.8.4 with default parameters [18]. Pairwise comparisons and clustering analysis were performed using Trinity v2.8.4 package (Biostars, New Taipei, Taiwan). Genes with at least 2-fold change with adjusted P < 0.05 were identified as differentially expressed genes (DEGs). Gene ID and Fragments Per Kilobase of transcript per Million mapped reads (FPKM) of genes in all the samples are listed in Supplementary Table S4. Gene Ontology (GO) enrichment analysis was performed by GENEONCOLOGY and PATRIC tools [19–22] and the chord plot was created by GO plot 1.0.2. Meanwhile, mechanisms of drugs that could kill *M. tuberculosis* were predicted and illustrated via the Kyoto Encyclopedia of Genes (KEGG) pathway database. In brief, significantly down-regulated genes in *M. bovis* BCG treated with antibiotics (logFC<-2 and FDR<0.01 compared with the DMSO control sample) that probably are critical for *M. tuberculosis* metabolism were selected and predicted as potential drugs of *M. tuberculosis* elimination.

### 2.9. Cell viability assay - LDH assay

The THP-1 monocytic cell line (ATCC TIB-202) was cultured in triplicate at density 5×10^4^ cells per well of a 96-well plate containing RMI1640 medium (GIBCO, USA) supplemented with 5% (v/v) fetal bovine serum (GIBCO, USA) at 37 °C with 5% CO2. The cells were then treated with 10-μl of sterile water and 10-μl of the compound (concentrations ranging from 0.2-μM to 200-μM), to determine spontaneous LDH activity (control group-one) and compound-treated LDH activity, respectively, using CyQUANT™ LDH Cytotoxicity Assay Kit (Thermo Fisher Scientific, USA). Additionally, cells of the control group -two were treated with nothing to measure LDH activity.

### 2.10. Statistical analysis

The differences in genes expression and *lux* signals among all samples were analysed using two-sample *t*-tests in GraphPad Prism (GraphPad Inc., USA). The standard of statistical significance was set at P-values ≤0.05.

## 3. Results

### 3.1. Construction and validation of the *lux*-based promoter-reporter platform

The *lux*-based *phoP* promoter-reporter plasmid (pMV306PhoP+Lux) and the negative control plasmid (pMV306Adaptor+Lux), were constructed as shown in Figure 1. *M. bovis* BCG transformed pMV306PhoP+Lux showed prominent *lux* when compared with negative control. Before the high throughput screening of small molecules, ETZ, a known inhibitor of PhoPR regulon via binding to *phoP* promoter, was used to validate if the *lux* signal of the reporter plasmid corresponded to the *phoP* expression in *M. bovis* BCG. The growth of *M. bovis* BCG (in terms of OD_600_) after ETZ treatment was also examined. The results showed that there was no significant difference in OD_600_ between *M. bovis* BCG with and without ETZ treatment (Figure 2a). However, the *lux* signal declined prominently upon ETZ treatment (Figure 2b). RT-qPCR also revealed that both expressions of *luxA* and *phoP* genes were significantly suppressed by 40% in the ETZ group (Figure 2c) when compared with the control group. The result assured that the *lux*-based *phoP* promoter-reporter plasmid responded to the chemical inhibitors in the same direction as *phoP* expression. The reduction in *lux* signal was unambiguously detectable using the 96-well plate reader.

**Fig. 2.**
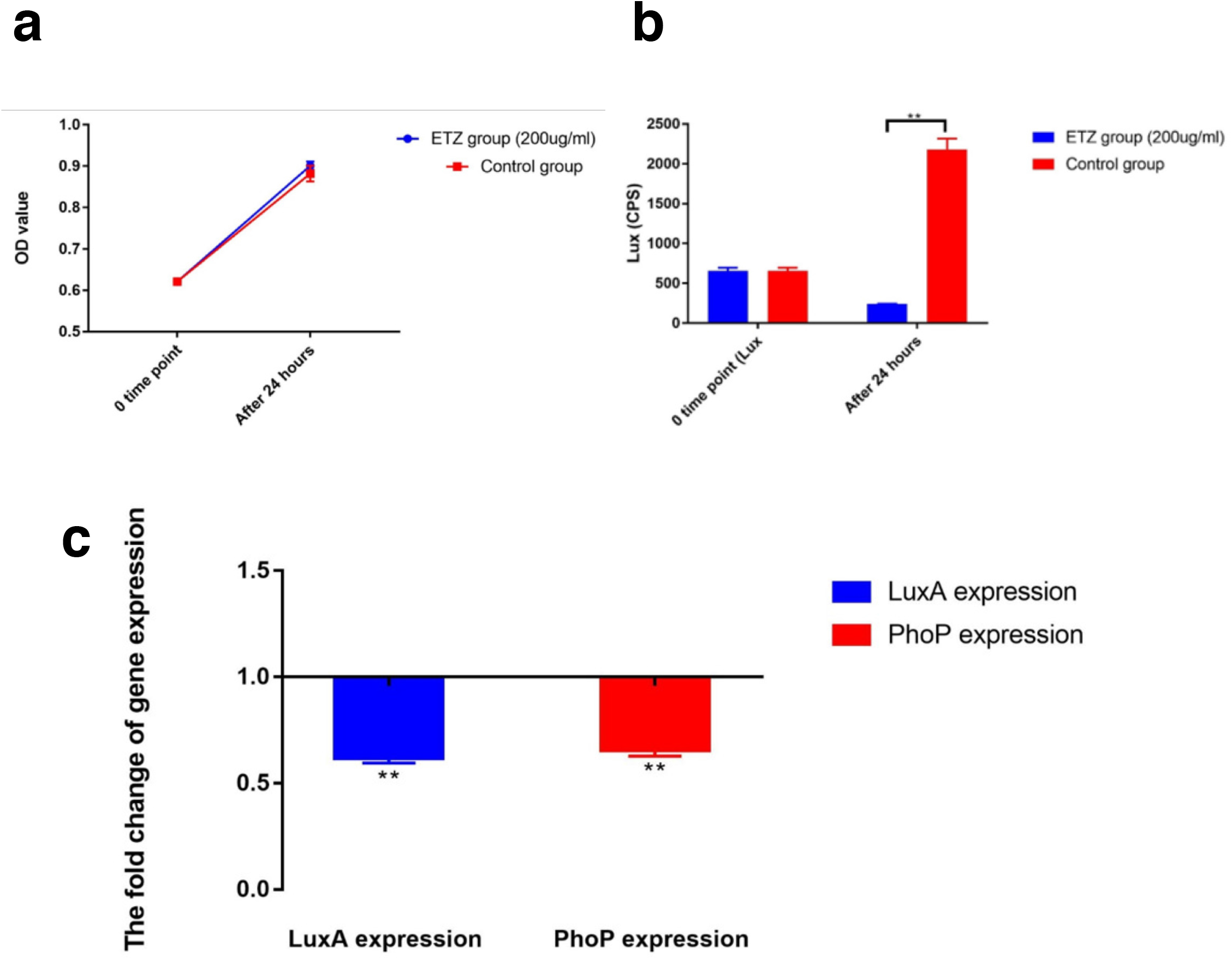
Validation of the *lux*-based promoter-reporter platform. **a,** OD_600_ in ETZ and control groups after 24-hour treatment. **b,** A LUX comparison between ETZ and control groups. **c,** *luxA* and *phoP* expressions of BCG Russia treated with ETZ based on a housekeeping gene *recX*.

### 3.2. Screening of 320 antiviral compounds and 3 anti-TB drugs

*M. bovis* BCG transformed with pMV306Adaptor+Lux (control set) and pMV306PhoP+Lux (phoP set) were respectively treated with a total of 320 antiviral compounds and 3 anti-TB drugs. The resulting *lux* signals and OD_600_ of each compound treatment were shown in Figure 3. For the control set, the *lux* signals were consistently below 50 CPS (no difference existed compared with the blank wells, *p*>0.05) after all compound treatments, except for triciribine (CPS =155) (Figure 3a). For the PhoP set, the drug-free controls (Figure 3b, green dots) showed an average *lux* signal at around 300 CPS, which is regarded as the basal *lux* intensity induced by the *phoP* promoter in the absence of compound challenges. When *M. bovis* BCG was treated with the three anti-TB drugs (Figure 3b, red dots), no significant changes of *lux* signals were observed. Notably, 16 antiviral small compounds (CPS ≈0, blue dots in Figure 3b) presented a significant suppression effect on *lux* signals after a 4-hour treatment when compared with the drug-free control (Figure 3c). These 16 compounds, which were listed in Supplementary Table S4, were selected for further experimental validation.

**Fig. 3.**
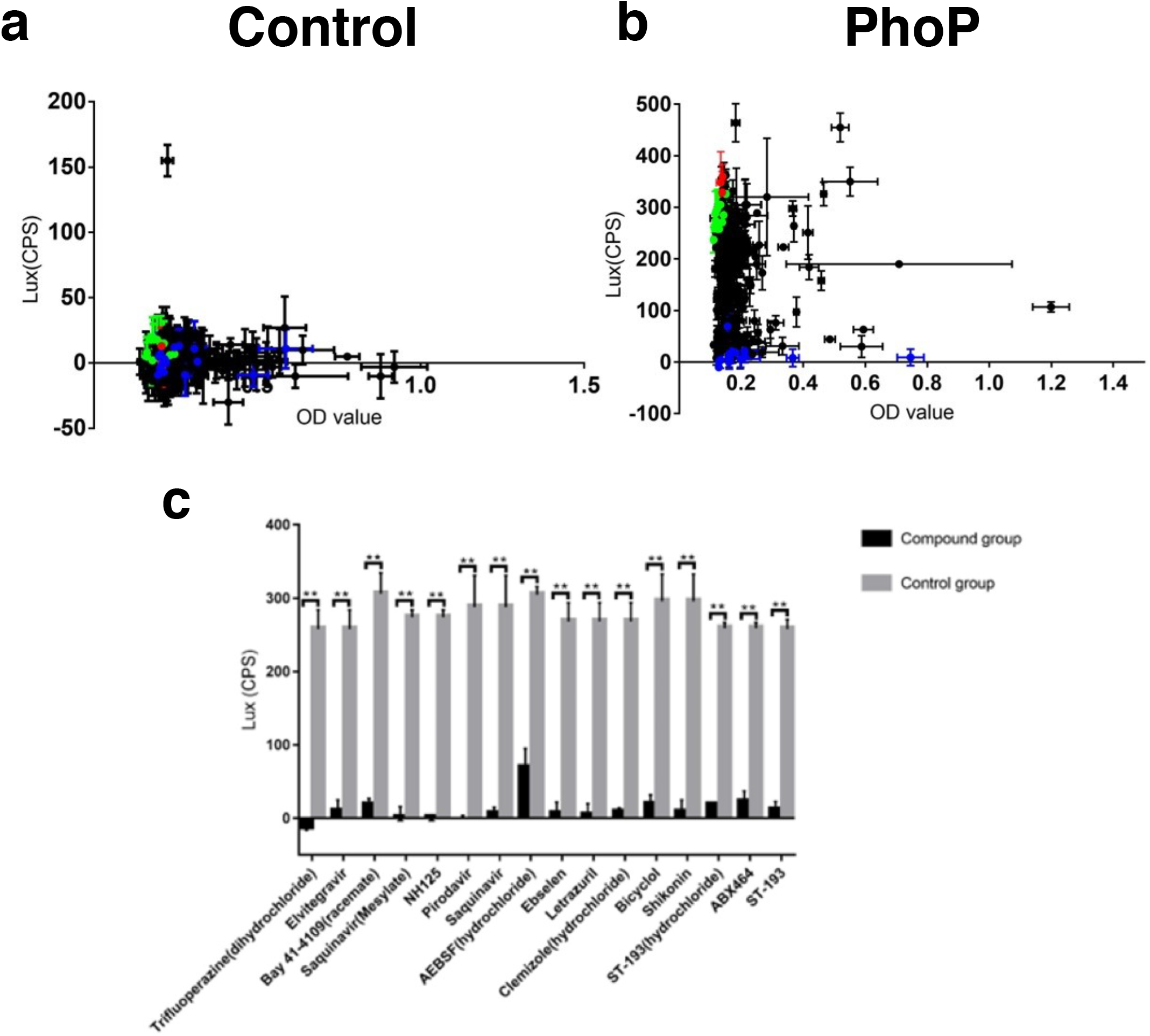
Screening of 320 antiviral compounds and 3 anti-TB drugs. **a**, OD_600_ and LUX of all samples with/without 4-hour compound treatment in negative control platform of BCG. **b**,, OD_600_ and LUX of all samples in *phoP* promoter-reporter platform. Green dots represent BCG samples (including pMV306Adaptor+Lux) without any compound treatment. Red dots represent samples treated with the three anti-TB drugs. Blue dots represent samples treated with the 16 selected compounds, while black dots represent samples treated with other compounds. **c**, Comparison of lux signals between compound treatment and control samples.

### 3.3. MICs and MBCs of six compounds against BCG/*M. tuberculosis*

*M. bovis* BCG after 4 hours of treatment with the above 16 compounds were washed and subcultivated in drug-free media, followed by incubation for 14 days. No viable *M. bovis* BCG was observed after treatment with six compounds, namely Trifluoperazine (dihydrochloride), Elvitegravir, NH125, Ebselen, Letrazuril, and Shikonin, whereas confluent growth was observed after treatment with the remaining 10 compounds. Subsequently, MICs and MBCs of the six compounds against *M. bovis* BCG, *M. tuberculosis* H37Rv, and two drug-resistant clinical isolates of *M. tuberculosis*, HKU14621 (MDR-TB) and WC274 (XDR-TB) were determined. Except for Elvitegravir and Letrazuril, these compounds demonstrated bactericidal effects on *M. bovis* BCG as well as all *M. tuberculosis* strains at a concentration of 100-μM or below (Supplementary Table S5). Notably, MICs and MBCs against MDR-TB and XDR-TB strains were almost identical to those against *M. tuberculosis* H37Rv, suggesting that these compounds can kill MDR- and XDR-*M. tuberculosis* clinical isolates as effective as the pan-susceptible reference strain (Supplementary Table S5).

### 3.4. RNA-Seq transcriptome analysis

To unveil the genetic mechanisms of drug actions of the 16 selected compounds on the *M. tuberculosis* complex, the transcriptomes of *M. bovis* BCG after compound treatment were profiled using RNA-seq. Based on the genome-wide differential expression patterns, the datasets could be clustered into two groups by principal component analysis (PCA) and hierarchical clustering (Figure 4A and 4B). Interestingly, the clustering separated the compounds with bactericidal activity from those without inhibitory effect. The compound group in which *M. bovis* BCG “survived” after the treatment was labelled as the survival (S) group, whereas the compounds that led to the “death” of *M. bovis* BCG were labelled as the dead (D) group. Interestingly, four compounds that belonged to the S group, namely Pirodavir, AEBSF (hydrochloride), Bicyclol and ABX464, exhibited similar transcriptome profiles as the D group. Only one gene, *phoY1*, was differentially expressed when compared to the compounds in the D group (Figure 5b, Supplementary Table S6A). These four compounds were therefore separated from the original S group and were assigned as S2 group (i.e., survival group with dead group-resembled transcriptomic profiles).

**Fig. 4.**
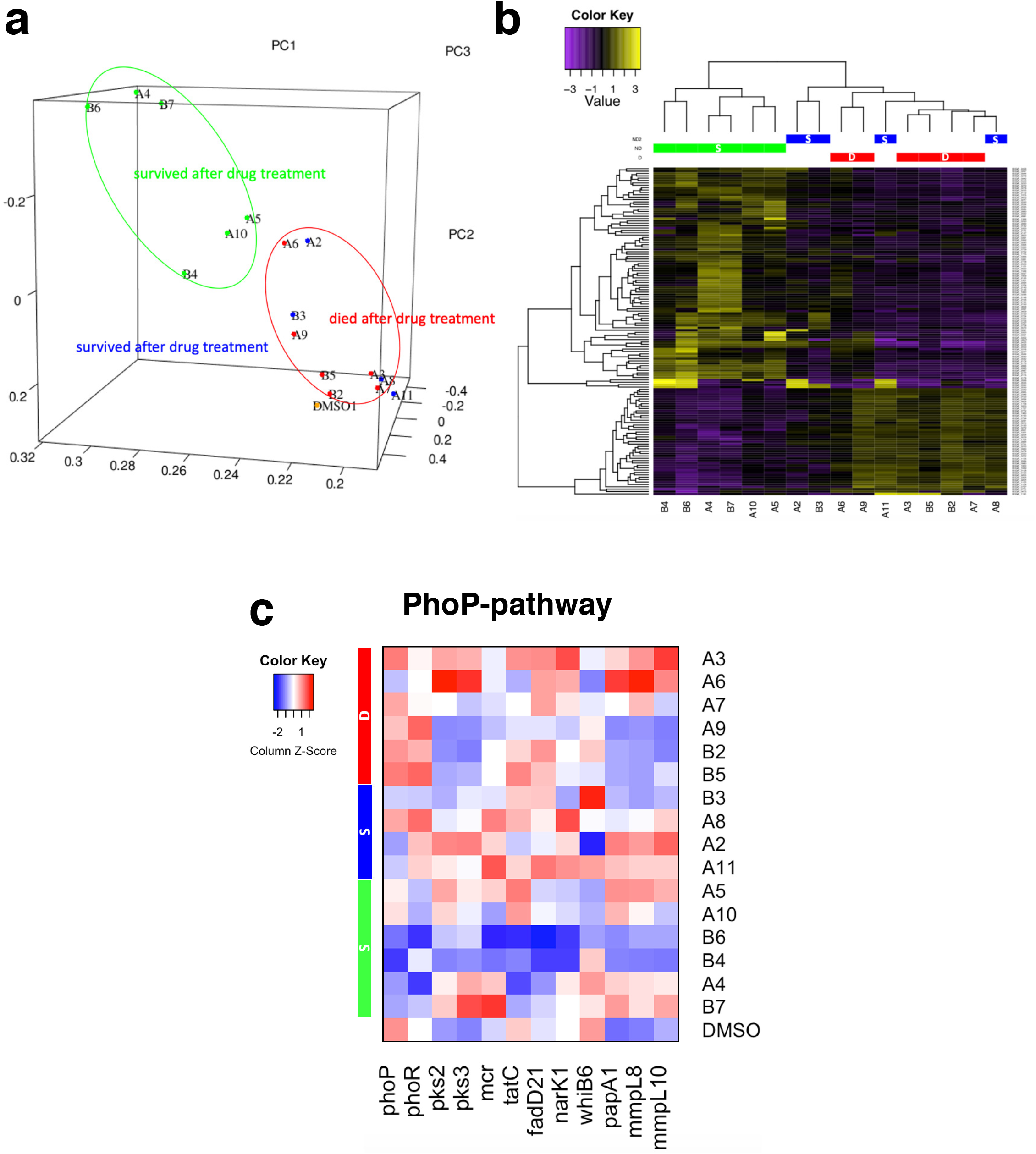
RNA-Seq transcriptome analysis and representative genes involved in phoP-pathway. **a**, Principal component analysis (PCA) on BCGs after treated with 16 compounds in 3 groups based on gene expression detected in all samples. **b**, Heatmap showing a clustering analysis of gene expression levels for the three groups of BCGs (*p*<0.05, FC>2 or <0.5). **c,** Expression of genes regulated by phoP/R in BCGs after treated with the 16 compounds.

**Fig. 5.**
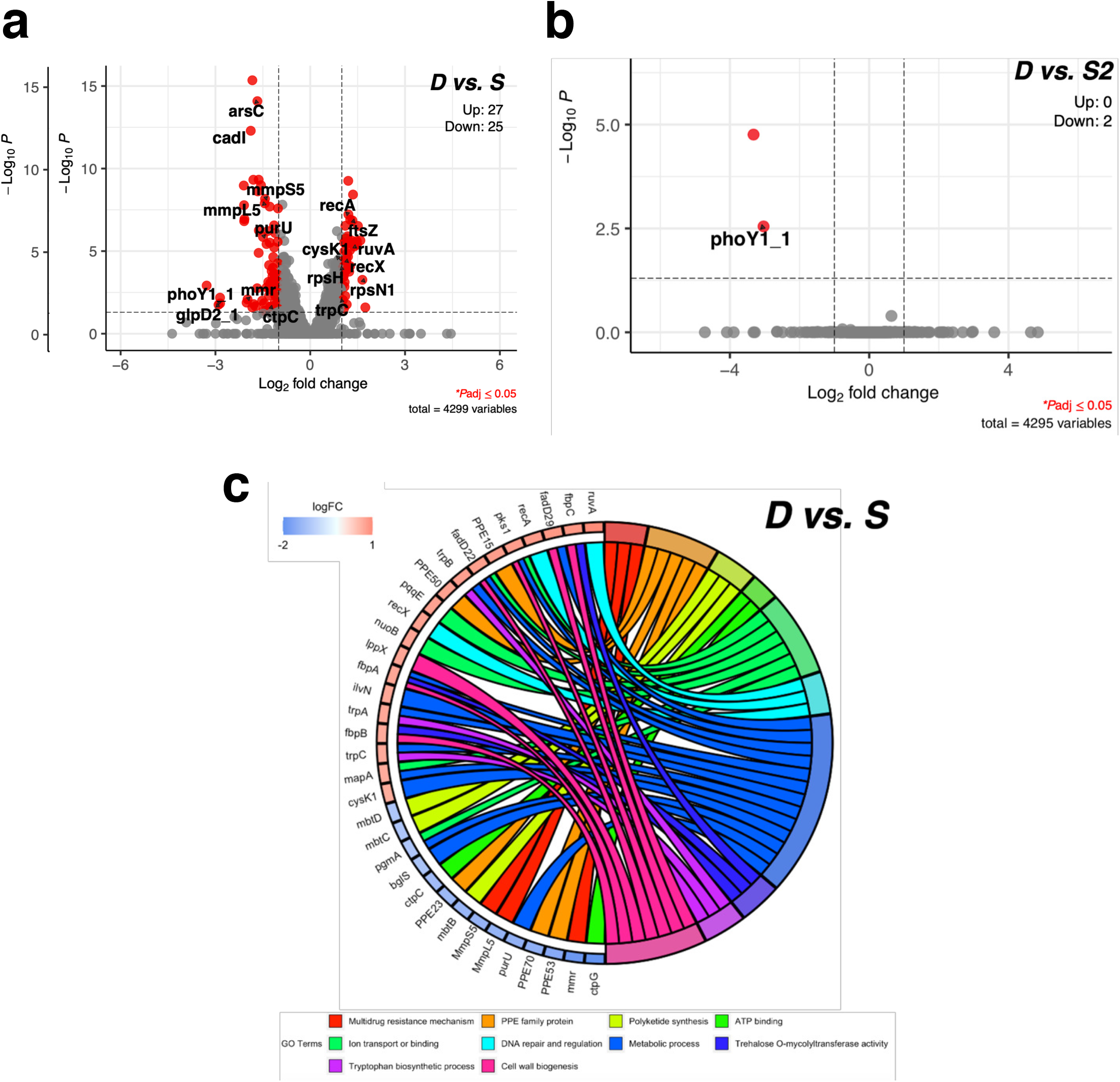
Molecular regulation associated with the anti-virulence process. **a**, Volcano plot DEGs between D and S. **b**, Volcano plot DEGs between D and S2. **c**, Molecular functions and biological processes enriched in DEGs for the survived groups.

To identify differentially expressed genes (DEGs) between the S and D groups, genes with at least 2-fold difference and adjusted P <0.05 were selected. Altogether, 52 DEGs (27 up-regulated and 25-down-regulated) were identified in groups D vs. S (Figure 5a, Supplementary Table 6B). Gene Ontology (GO) enrichment analysis was performed to reveal important biological processes and molecular functions dysregulated in the two groups. We demonstrated that genes involved in multidrug resistance mechanisms (eff*lux* pump), DNA repair system, PPE family protein and polyketide synthesis were down-regulated in the dead group while genes related to the DNA repair process and cell wall biogenesis were down-regulated in the survival (S) group (Figure 5C, Supplementary Table S7A). The results highlighted the importance of these processes during anti-virulence.

### 3.5. Expression of *phoP*-associated pathways upon compound treatment

The expression of *phoP* and its downstream gene network was investigated specifically. Surprisingly, *phoP* expression was up-regulated in response to the challenges by the compounds in the dead (D) group. Conversely, the expression of *phoP* was mostly suppressed in the survival group (S group). Notably, in addition to *phoP*, the related genes in the associated pathways were consistently down-regulated when *M. bovis* BCG was treated with two compounds, ST-193 (B4) and ST-193 (hydrochloride) (B6) (Figure 4c). This implicated that these two compounds were likely to dysregulate the entire *phoP*-associated pathways.

### 3.6. Prediction of drug targets in the dead group

As described above, six compounds exhibited bactericidal activity against *M. bovis* BCG. The DEG profiles were investigated to identify the potential drug targets of these compounds. Some common down-regulated genes (logFC<-2 and FDR<0.1 when compared with the DMSO control sample), including *cysA2, frdC*, and *glpD2_2*, were identified when *M. bovis* BCG was treated with Ebselen (A3), NH125 (A9) and Shikonin (B2) (Supplementary Table S7B). It should be noted that the sequences of *cysA2, frdC*, and *glpD2_2* in *M. bovis* BCG were almost 100% identical to those in *M. tuberculosis* H37Rv. It is consistent with our observation in phenotypic drug susceptibility tests that Ebselen, NH125 and Shikonin shared very similar MICs and MBCs against *M. bovis* BCG and *M. tuberculosis* strains. Therefore, *cysA2*, *frdC*, and *glpD2* were considered as potential drug targets of these small compounds.

### 3.7. THP-1 cytotoxicity of the small-molecule compounds

The cytotoxicity of the 16 selected small compounds was determined using THP-1 macrophage. Overall, five compounds were considered as non-cytotoxic to human macrophage given that their CC_50_ were greater than 200-μM, the highest achievable concentration in this study (Figure 6, Supplementary Table S8). Of these five compounds, ST-193 and ST-193 (hydrochloride) were shown to dysregulate the expression of entire *phoP*-associated pathways without inhibiting the *in vitro* growth of *M. bovis* BCG, suggesting that they were potential anti-virulence agents in anti-TB therapy. Moreover, among the compounds that had a bactericidal effect on *M. tuberculosis*, only Ebselen had the CC_50_ (>200-μM) greater than its MIC (50-μM) and MBC (100-μM) against the *M. tuberculosis* complex. It was therefore considered as a potential antibiotic for *M. tuberculosis*.

**Fig. 6.**
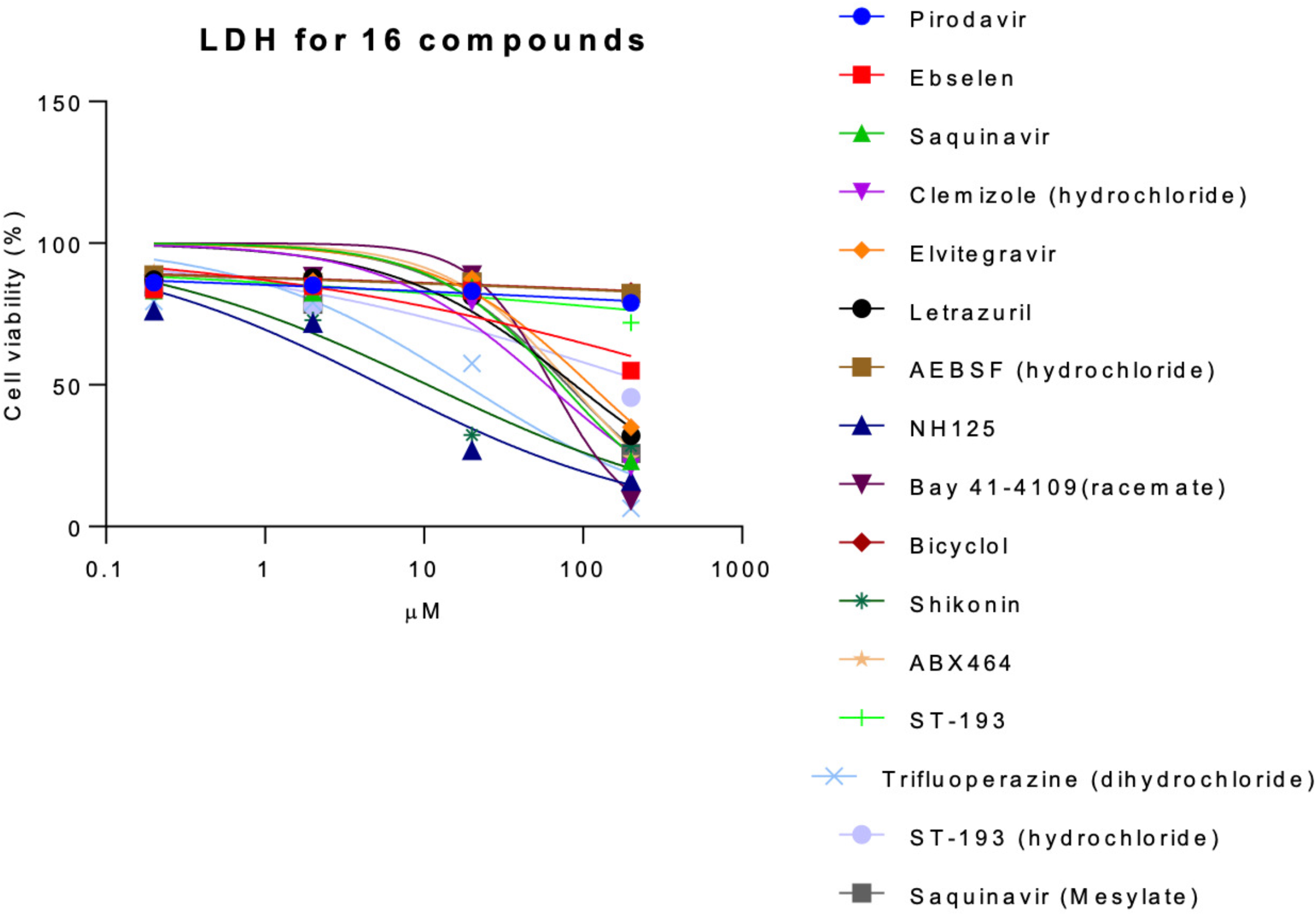
LDH results of 16 compounds in THP-1 cells.

## 4. Discussion

Repurposing bioactive small-molecule compounds have been suggested as a new therapeutic approach against *M. tuberculosis* infection. Unlike previous studies, which focused on one or several compounds [8], we examined the anti-TB potency of a library of 320 small-molecule compounds with antiviral activity. As antiviral agents, these compounds are expected to have good penetration across host cell members, which is an important feature of the drugs used to treat intracellular pathogens, such as *M. tuberculosis* complex.

In addition to identifying new candidates of antibiotic, this study aimed to discover anti-virulence agents, which impaired the bacterial virulence factors without killing the organisms, for *M. tuberculosis* [7]. Notably, the development of anti-virulence drugs requires an in-depth understanding of the roles that diverse virulence factors have in disease processes. Previous studies from our team and other research groups suggested that *phoP* is essential for the intracellular growth of the *M. tuberculosis* complex inside macrophages [10, 11]. Disruption of *phoP* could impair intramacrophage and facilitate the host’s immune system to eradicate the bacteria.

Instead of individual measurement of *phoP* expression using RT-qPCR, we developed a high throughput *lux*-based promoter-reporter screening platform to select potential agents that can inhibit *phoP* promoter activity. This platform enabled real-time measurement of *phoP* expression in viable *M. bovis BCG* culture in response to the compound challenge, in terms of *lux* signal, in a batch of 96 samples. In the screening platform validation, we successfully demonstrated that ETZ could reduce both the RNA quantity of *phoP* and *lux* signals, indicating the luminescence generated by the phoP promoter-reporter plasmid, pMV306PhoP+Lux, corresponded to *phoP* expression in *M. BCG* bovis.

By using the high throughput screening platform, 16 out of 320 (5.0%) small-molecule compounds were identified as potential candidates against *M. tuberculosis* infection. Surprisingly, six of them: trifluoperazine (dihydrochloride), elvitegravir, NH125, ebselen, letrazuril, and shikonin did not show a decrease in *phoP* gene expression and directly eliminated *M. bovis* BCG. The decrease in their *lux* signals was possibly caused by bacterial death. In the molecular regulation analysis, we highlighted that dysregulation of multidrug resistance mechanisms, DNA repair system, PPE family protein and polyketide synthesis would determine *M. tuberculosis* death. Among the six compounds, only Ebselen exhibited CC_50_ greater than its MIC and MBC against the *M. tuberculosis* complex. Our RNA-seq analysis identified that Ebselen displayed bactericidal activity in *M. bovis* BCG through *cysA2* and *glpD2* down-regulation which are the major regulators in sulfur and glycerol metabolism respectively. *CysA2* is an essential regulator in the sulfur assimilation pathway [23] and its dysregulation could impair *M. tuberculosis* survival in macrophages [24]. In parallel, a previous study demonstrated that inhibition of glycerol metabolism by 2-aminoquinazolinones via *glpD2* down-regulation could kill *M. tuberculosis in vitro* [25]. Taken together, dysfunctions of sulfur and glycerol metabolism are possible drug mechanisms for Ebselen. It might also be effective on *M. tuberculosis* as its sequences of *cysA2* and *glpD2* were almost 100% identical to those in *M. bovis* BCG. Hence, Ebselen could be considered as a potential antibiotic for the *M. tuberculosis* complex.

Among the remaining 10 survival candidates, six of them: Saquinavir, Clemizole (hydrochloride), Bay 41-4109 (racemate), ST-193, ST-193 (hydrochloride) and Saquinavir (Mesylate) exhibited a distinct transcriptome profile and lower gene expression of *phoP*. In the molecular regulation analysis, genes related to the DNA repair process and cell wall biogenesis in this group were significantly down-regulated. It was found that the activity of the PhoP-related system interacted with genes involved in the fadD family which regulates the fatty acid β-oxidative pathway [26]. Our results revealed that genes in the fadD family (*fadD29* and *fadD22*) are of lower expression in *M. bovis* BCG after being treated with these six compounds. Disruption of fatty acid biosynthesis could inhibit bacterial differentiation and growth [27]. These six compounds are potential anti-virulence agents because they inactivated *M. bovis* BCG through DNA damage, cell-wall destruction and differentiation inhibition, thereby facilitating the host’s immune system to eradicate the bacteria. Interestingly, ST-193 and ST-193 (hydrochloride) of CC_50_ greater than 200-μM dysregulated the entire *phoP*-associated gene network in *M. bovis* BCG so they are the most desirable anti-virulence agents.

For proof-of-concept, only 320 small compounds were screened in this study. Despite this small-scale screening, three (0.94%) compounds were identified as potential antibiotic and anti-virulence agents *against M. tuberculosis*. In the future, the discovery of more new candidates of anti-TB drugs is anticipated if more drug repurposing compounds are screened using the promoter-reporter platforms coupled with automatic high throughput screening instruments. Current TB treatment relies on a synergistic combination of drugs administered for the desired time to ensure definitive non-relapsing cures and avoid drug-resistant mutants [28]. In 2021, the World Health Organization (WHO) established a position statement on “*Position statement on innovative clinical trial design for development of new TB treatments*” to outline clinical trial designs [29]. Anti-virulence agents facilitate the development of new TB therapies and provide us an avenue to establish our innovative TB treatment in clinical trials. However, it is necessary to see if the anti-virulence agents have synergistic or antagonistic interaction with existing anti-TB drugs. Fractional inhibitory concentrations of the compounds and the current anti-TB drugs should be determined by checkerboard assay in future studies.

## 6. Conclusion

In this study, a *lux*-based promoter-reporter platform was constructed to screen a total of 320 small compounds for anti-TB potency. Eventually, Ebselen was identified as the most desirable antibiotic, while ST-193 and ST-193 (hydrochloride) were was potential candidates of anti-virulence for the *M. tuberculosis* complex.

## Author Contributions

ZL, AL, PG, RK and GS conceived the original concept and designed the study. ZL, KW, KT, and RR performed the experiments. ZL and AL analysed the data and wrote the original paper. GC, TN, CC, HL and GS amended and revised the writing. All authors read and approved the final version of the manuscript.

